# Disentangling the aging network of a termite queen

**DOI:** 10.1101/2020.12.19.423576

**Authors:** José Manuel Monroy Kuhn, Karen Meusemann, Judith Korb

## Abstract

**Background:** Most insects are relatively short-lived, with a maximum lifespan of a few weeks, like the aging model organism, the fruit-fly *Drosophila melanogaster*. By contrast, the queens of social insects (termites, ants, some bees and wasps) can live for more than a decade. This makes social insects promising new models in aging research providing insights into how a long reproductive life can be achieved. Yet, aging studies on social insect reproductives are hampered by a lack of quantitative data on age-dependent survival and time series analyses that cover the whole lifespan of such long-lived individuals. We studied aging in queens of the drywood termite *Cryptotermes secundus* by determining survival probabilities over more than 15 years and performed transcriptome analyses for queens of known age that covered their whole lifespan.

**Results:** The maximum lifespan of *C. secundus* queens was 13 years with a median maximum longevity of 11.0 years. Time course and co-expression network analyses of gene expression patterns over time indicated a non-gradual aging pattern. It was characterized by networks of genes that became differentially expressed only late in life, namely after an age of 10 years, which associates well with the median maximum lifespan for queens. These old-age gene networks reflect processes of physiological upheaval. We detected strong signs of stress, decline, defence and repair at the transcriptional level of epigenetic control as well as at the post-transcriptional level with changes in transposable element activity and the proteostasis network. The latter depicts an upregulation of protein degradation, together with protein synthesis and protein folding, processes which are often down-regulated in old animals. The simultaneous upregulation of protein synthesis and autophagy is indicative for a stress-response mediated by the transcription factor *cnc*, a homolog of human *nrf* genes.

**Conclusion:** Our results show non-linear senescence with a rather sudden physiological upheaval at old-age. Most importantly, they point to a re-wiring in the proteostatis network as central for explaining the long life of social insect queens.

## Background

Almost all animals age, but at different pace [1]. The fruit fly *Drosophila melanogaster* lives only for around seven weeks [2], while the clam Ocean Quahog, *Arctica islandica*, can have a lifespan of more than 400 years [3]. Generally, organisms with large differences in rates of aging are found between widely divergent species [1,4], which makes controlled comparisons of the underlying aging mechanisms difficult.

Classical model organisms typically have a short lifespan and can be characterized by r-life history strategies (‘live fast, have many offspring and die young’) as exactly these traits make them good model organisms. Social insects such as termites, ants, or the honeybee, offer promising new insights into aging research because individuals with the same genetic background can differ by orders of magnitudes in lifespan. Within a social insect colony, which is generally a large family, the reproducing queen (and in termites, also the king) can reach lifespans of more than 20 years, while non-reproducing workers have a lifespan of a few months only [5–8]. However, quantitative demographic data covering the whole lifespan of queens are inherently rare and many reports on queen-longevity are more anecdotal. Thus, it is largely unknown for long-lived queens whether they age gradually or whether aging is a more sudden event.

During recent years, several pioneering studies, especially on the honeybee, revealed exciting new insights into the mechanisms of how queens can live so long. In the honeybee, juvenile hormone (JH) seems to have lost its direct gonadotropic function in adults so that queens have a high expression of the yolk precursor gene vitellogenin (*Vg*) without requiring high JH titres (*e.g.*, [9,10]). This result has led to the hypothesis that uncoupling JH and *Vg* expression might account for the long life of honeybee queens [9] as well as social insect queens more generally [11] because the life-time shortening consequences of high JH titers are absent. However, this re-wiring along the JH-*V_g_* axis is not universal for all social Hymenoptera, since the queens of many other ant and bee species require JH for vitellogenesis (*e.g.* [12] and references therein). For termites, fewer studies exist, but JH is required for vitellogenesis [13,14]. Hence, other mechanisms must exist to explain the long life of termite queens. Studies of the subterranean termite *Reticulitermes speratus* implicated the involvement of a *breast cancer type 1 susceptibility (BRCA1*) homolog [15], which is involved in DNA repair [16], and better protection against oxidative stress by superoxide dismutases and catalases [17,18]. The latter have also been discussed for other social Hymenoptera, including ants and the honeybee. Yet overall evidence of the role of oxidative stress is less clear (reviewed in [19–21]). Furthermore, regulation of the activity of transposable elements (TEs) [22], and changes in the insulin/insulin-like growth factor1 signalling (IIS) and target of rapamycin (TOR) pathways [23] have been linked with caste-specific aging differences in termites. Both, the TOR and IIS pathway, have been associated with longevity in model organisms from *D. melanogaster* to mice and humans [24–26]. They are the most intensively studied aging related pathways and they have also been associated with caste differences in social Hymenoptera (*e.g.,* [27–31]). Yet, all studies on social insects suffer from a lack of time-series data to investigate molecular changes across the lifespan of long-lived queens. Like the demographic life history data, such data are inherently difficult to obtain due to the long lifespan of queens. However, they are necessary (i) to understand the aging process, (ii) to work out potential changes compared to solitary insects, and (iii) to identify the relevant age-classes for detailed studies. The latter is a completely overlooked issue but highly relevant. If, for example, aging is a non-linear process, differences across studies might just be consequences of none-comparable age-classes between studies.

We studied aging in termite queens of known age across their entire lifespan to measure at the ultimate, eco-evolutionary level age-dependent survival and at the proximate, mechanistic level age-specific changes in gene expression. For the latter, we generated head and thorax transcriptomes of queens of different age (for an outline of the workflow see Additional File 1, Figure S1). We used field collected, newly established colonies of the wood-dwelling termite *Cryptotermes secundus* (Hill, 1925) (Blattodea, Isoptera, Kalotermitidae) that were kept under identical conditions in the laboratory for a time span of 15 years. Keeping them under constant and optimal conditions allowed us to study intrinsic aging, disentangled from causes of extrinsic mortality such as predation, food shortage, or disease. As typical for wood-dwelling termite species, *C. secundus* colonies are founded in a piece of wood, which serves as food and shelter and which workers never leave to forage outside. Such species have a low social complexity with small colonies and totipotent workers that develop into sexuals.

## Results

### Survival analysis

Overall, we had surviving founding queens from an age of two years up to a maximum of 13 years. The potentially 14- and 15-year old queens all had died as had most queens with a potential age of ≥ 12 years. Out of eight queens in this ‘old-age’ class, only a single queen (13 years) had survived (Fig. 1). The median longevity of the queens in the laboratory after successful colony establishment was estimated with Kaplan Meier survival analysis to be 12.0 years (SE: ± 0.54) (mean longevity: 11.1 years, SE: ± 0.66) (Fig. 1).

**Figure 1.**
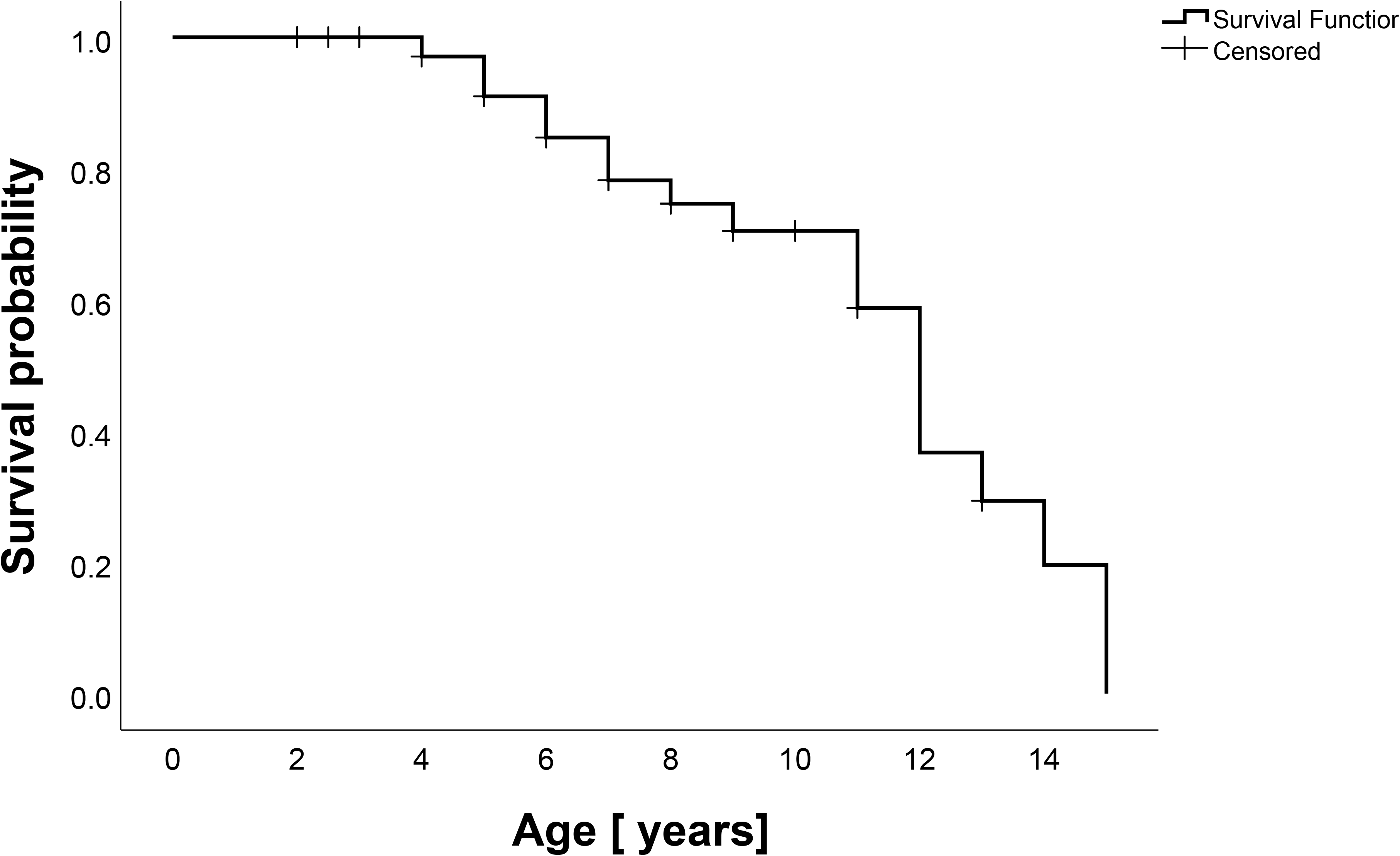
Survival plot of *C. secundus* queens. Shown is the age-dependent survival probability of queens. The median longevity of queens in the laboratory after successful colony establishment was estimated with Kaplan Meier survival analysis to be 12 years (mean longevity: 11.1 years). The maximum lifespan was 13 years. After an age of around 11 years, life expectancy declines rapidly; out of eight queens with a potential age ≥ 12 years, all had died, except one 13-year old queen.

### Identifying transcripts that change their expression with age: age-related DETs

To study gene expression changes over the lifetime of queens we generated transcriptomes of head and thorax from twelve queens with different chronological age since onset of reproduction, from two until 13 years, covering the complete lifespan of *C. secundus* queens: 2, 3, 4, 5, 6, 7, 8, 9, 10 (two samples), 11, and 13 years (Additional File 2, Table S1).

A total of 169 transcripts were significantly differentially expressed (DETs) over time as revealed by Iso-MaSigPro time series analysis (Additional File 2, Table S2). According to their expression pattern, DETs were grouped into six Iso-MaSigPro clusters (hereafter, ‘cluster’) (Fig. 2). Cluster 1 represented 44 DETs, which were slightly expressed in young queens followed by a decline at middle ages and a strong increase when queens became older. The 32 DETs of cluster 2 characterized young queens with a declining expression with age. Clusters 3 and 5 comprised 31 and 37 DETs, respectively, that were highly expressed in middle aged queens, while cluster 4 and cluster 6 (15 and 10 DETs) characterized old queens with no expression in young ones. Thus, in the following we referred to the DETs as young (cluster two), middle-aged (clusters three and five) and old DETs (clusters one, four and six). Details for all clusters are provided in Additional File 2, Table S2.

**Figure 2.**
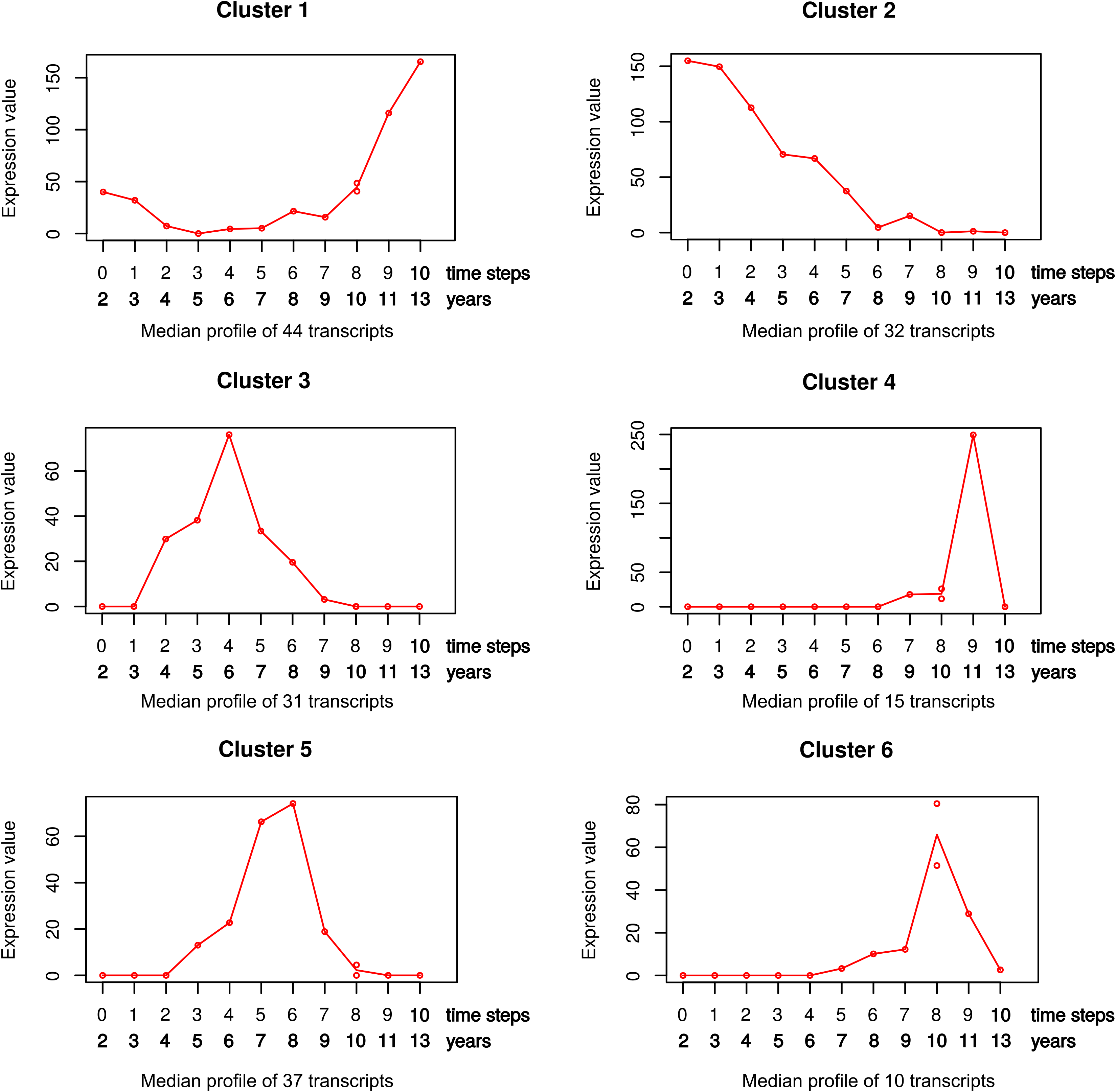
Median expression profiles of DETs assigned to Iso-MaSigPro clusters. Iso-MaSigPro grouped the differentially expressed transcripts (DETs) into six clusters. DETs of cluster 1, 4 and 6 were especially highly expressed in old queens, while those of cluster 3 and 5 characterized middle-aged queens and those of cluster 2 young queens. The expression values correspond to normalized counts (see Methods). The youngest queen (age: 2 years) was taken as time step zero and each of the subsequent older queens (based on chronological age) were considered to be one time step older. One age class (time step 8; age: 10 years) consisted of two samples.

### Identifying modules of co-expressed transcripts

To identify modules of co-expressed transcripts, we performed a weighted gene co-expression network analysis (WGCNA). It revealed a total of 254 modules of co-expressed transcripts. Based on eigengene values, 13 modules correlated significantly positively with age and 13 negatively (see Additional File 1, Figures S2 and S3; Additional File 3 (WGCNA module-age association, shown are eigengene values for all modules).

### Identifying transcript co-expression modules with age-related DETs

Within the age-correlated WGCNA modules, we identified age-related DETs. The negatively age-correlated module ‘seashell4’ had the highest number of young DETs (10 DETs). No gene ontology (GO) term was enriched for this module. The highest number of old DETs were found in the positively age-correlated modules ‘cyan’ (89 DETs) and ‘tan’ (79 DETs) (Additional File 2, Table S3 and S4). Only broad categories were enriched in the ‘cyan’ module (*e.g.* RNA metabolic process and gene expression) and the ‘tan’ module was enriched for ribosomal and tRNA related functions (Additional File 1, Figure S4).

### Extracting age-related subnetworks based on age-related DETs

To generate subnetworks related to the age-related DETs, we located them in the WGCNA co-expression network. These DETs and their one- and two-step neighbors (*i.e.,* the ‘second level neighborhood’) were then extracted from the co-expression network, which resulted in 50 subnetworks of different sizes (for more details see Methods) (Additional File 1, Figure S5). Note, DETs might be located at the boundaries of multiple WGCNA modules which means the subnetworks obtained consist of fragments of multiple WGCNA modules. The resulting subnetworks either contained young and middle-aged DETs or old DETs, with a single exception where a middle-aged DET was in the periphery of the largest subnetwork containing old DETs. The largest subnetwork containing either young and middle-aged DETs (hereafter, young subnetwork) or old DETs (plus a single middle-age DET; hereafter old subnetwork) were further analyzed.

#### Young subnetwork

The largest young subnetwork comprised 164 transcripts (out of these 24 Iso-MaSigPro DETs) of which only 12 (7%) were one-to-one orthologs to *D. melanogaster* genes (Additional File 2, Table S3). The GO enrichment analysis of the young subnetwork showed multiple Biological Process (BP) terms related to RNA catabolism, but these GO terms were not significant after correcting for multiple testing (Additional File 2, Figure S6).

##### TE activity and genome instability

53 DETs (32%) of the young subnetwork were related to TEs (Fig. 3 and Additional File 2, Table S3), comprising TEs and genes from TE defense pathways. This included one homolog of the gene *argonaute 2* (*ago2*) (two transcripts), an essential gene of the endo-siRNA pathway which silences TEs [32], and *arsenite 2* (*ars-2*) which is required for siRNA and miRNA-mediated TE silencing [33]. Additionally, we found two genes connected to DNA damage response and genome instability: *kin17* and *PIF1-like* gene.

**Figure 3.**
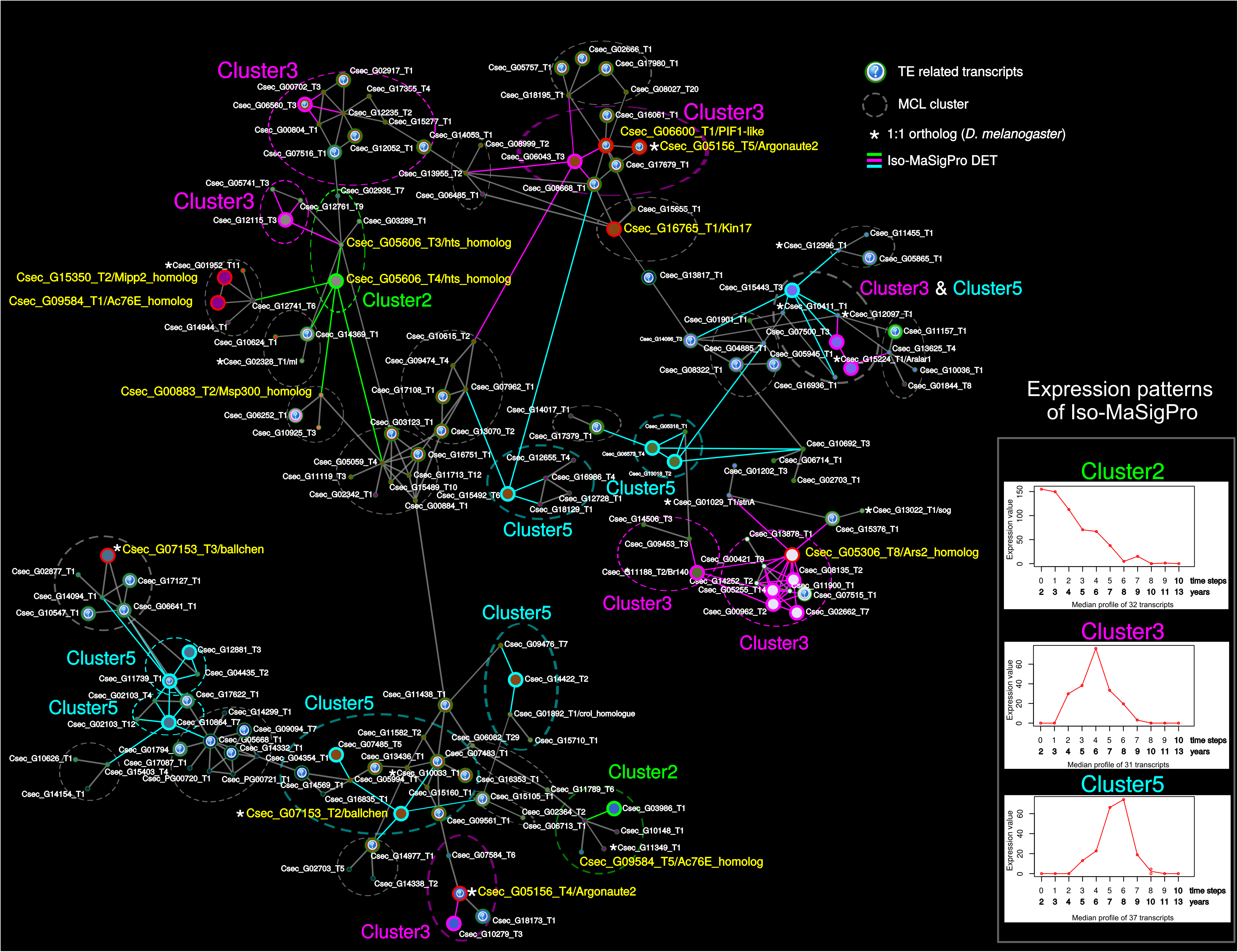
Young subnetwork highlighting Iso-MaSigPro DETs. Shown is WGCNA-based co-expression network of transcripts, which contains DETs characterising young and middle-aged queens and their one- and two-step neighbors (*i.e.*, young subnetwork; for more information, see text and Additional File 1, Figure S1 and S2). Highlighted are the Iso-MaSigPro DETs of cluster 2, 3, and 5, characterizing young and middle-aged queens (see insert; Fig. 2). Node colors correspond to the WGCNA modules. Transposable element (TE) related transcripts are highlighted with a ‘?’. Transcripts with an asterisk indicate 1:1 orthologs (*C. secundus* and *D. melanogaster*). Connection length and width do not have a meaning. Red circles indicate transcripts discussed in the text.

##### Other signatures

From well-known aging pathways, we identified (i) *inositol polyphosphate phosphatase 2* (*mipp2*) and (ii) *adenylyl cyclase 76E* (*ac76E*). The former is part of the TOR pathway and has been associated with longevity [34] and the latter is activated by the transcription factor ‘Forkhead box O’ (*foxo).* Additionally, we found several fecundity related DETs. They included two transcripts of the gene *hu li tai shao* (*hts*) (one a DET of IsoMaSigPro cluster two) and one homolog of the gene *bällchen* (*ball*) (two transcripts) (one a DET of cluster five, Fig. 3)

#### Old subnetwork

The largest ‘old subnetwork’ comprised 1,098 transcripts (out of these 42 Iso-MaSigPro DETs). 521 transcripts (47%) were identified as one-to-one orthologs of *D. melanogaster* genes (Additional File 2, Table S4). Iso-MaSigPro DETs in the old subnetwork belonged mainly to Iso-MaSigPro clusters 1 and 4. The second level neighborhoods of these DETs were connected in the network and a GO enrichment analysis revealed multiple GO terms associated with protein related functions, including translation, protein folding, unfolded protein binding, proteolysis involved in cellular protein catabolic process, protein targeting to ER, ribosome, and proteasome complex (Additional File 1, Figures S7 and S8). 198 transcripts of the old subnetwork (18%) were involved in protein translation, protein folding, and protein catabolism and proteolysis (Figs. 4 and 5 [35,36] and Additional File 2, Table S4). Additionally, 61 transcripts (~6%) were related to TEs (Additional File 2, Table S4).

**Figure 4.**
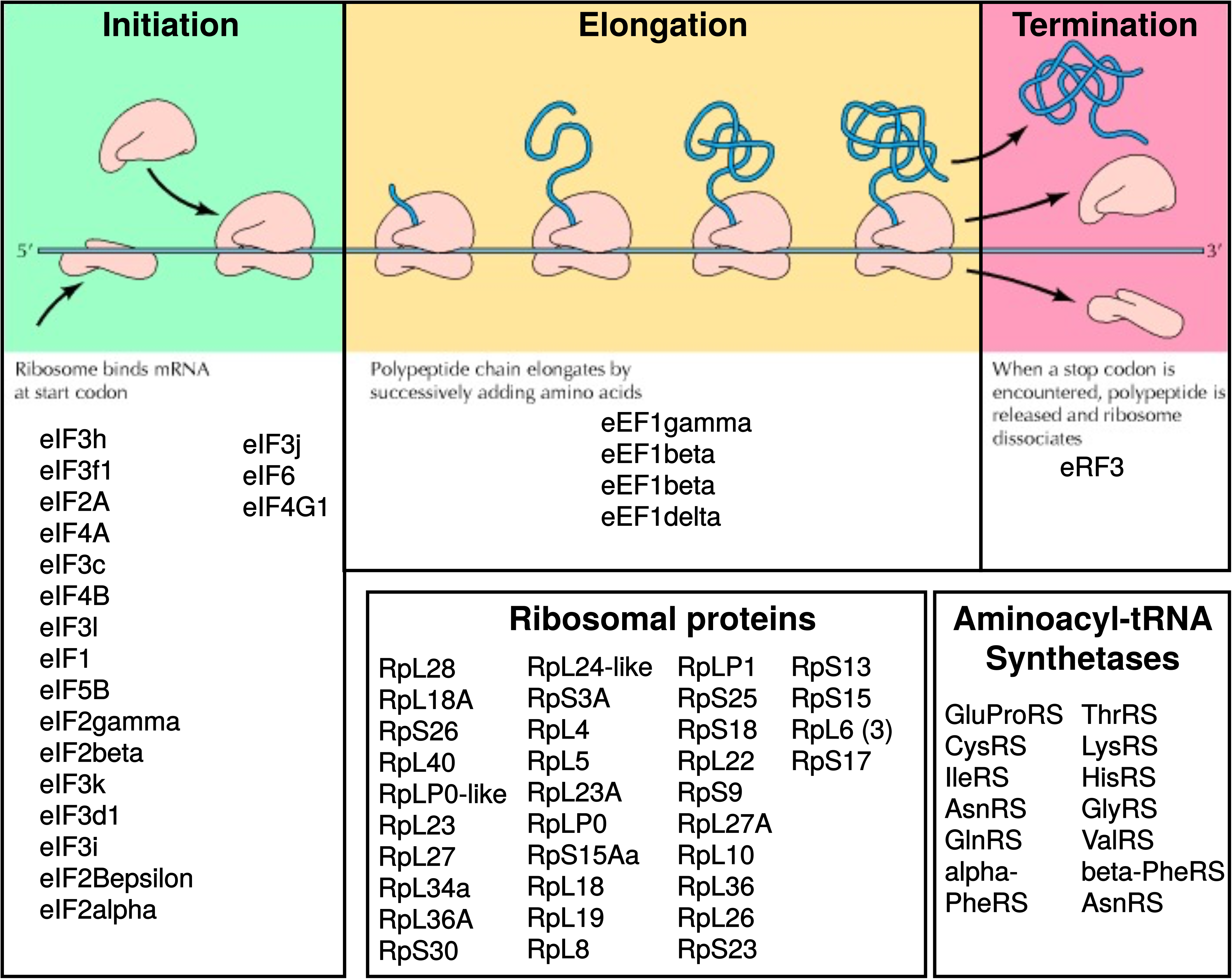
Genes related to protein synthesis that were found in the old subnetwork. Shown are genes that have been related to various processes of protein synthesis, from initiation, and elongation to termination. For all genes listed, corresponding transcripts were present in the old subnetwork of *C. secundus* queens. Figure modified after [35].

**Figure 5.**
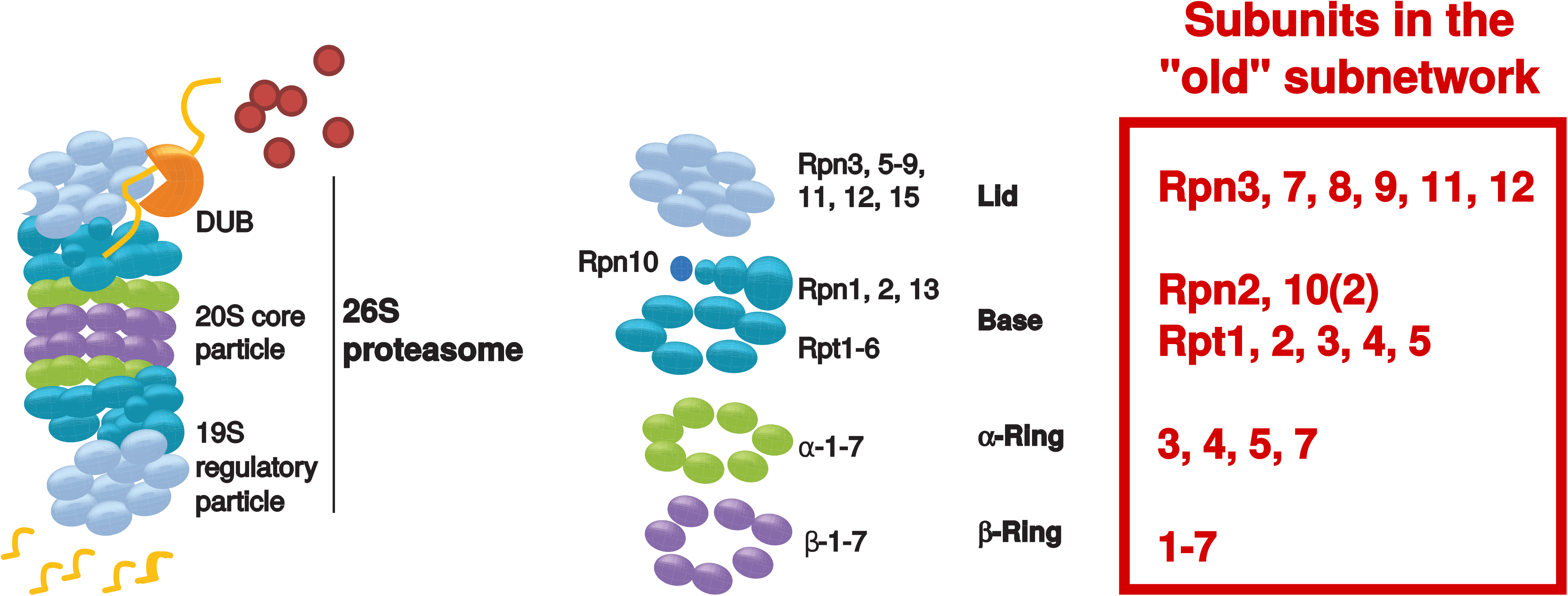
Genes related to the proteasome complex that were found in the old subnetwork. Shown are genes that have been related to the proteasome complex. The textbox in red indicates subunits, for which we found transcripts in the old subnetwork. Figure modified after [36].

##### Epigenetic modifications, transcriptional regulation, and TE activity

Many genes of the old subnetwork are involved in de/acetylation and methylation of DNA, which are important epigenetic modifications that regulate gene expression and genome stability [37–39] (Additional File 2, Table S4).

Most strikingly, two crucial histone acetylation modifying complexes, the Tip60 acetyltransferase complex and the male specific lethal (msl) complex were represented in the old subnetwork. The former included the genes *dom, ing3, mrg15, pont* and *rept,* and the latter *msl-1*, *msl-3* and *mof*. Genes involved in deacetylation of DNA were, for instance, *sirtuin 1* (*sirt1*), *histone deacetylase 3* (*HDAC3*), and *histone deacetylase 6* (*HDAC6*). Genes linked to epigenetic histone methylation included, for instance, *ash-1* and *lid*. Another well represented group of genes connected to expression regulation in the old subnetwork were spliceosome components and splicing factors. Additionally, we found in the old subnetwork important transcripts related to TE silencing: *dicer-2*, *Hsc70-4, Hsc70-3, Hsp83, trsn*, *armi, Rm62, Gasz, Tudor-SN,* and *Hel25E.* Details are given in Additional File 2, Table S4.

##### Proteostasis and oxidative stress

Related to proteostasis we detected a strong signal for protein synthesis and degradation. Regarding protein synthesis, the old subnetwork comprised many transcripts coding for initiation, elongation and termination factors, as well as many ribosomal proteins and aminoacyl-tRNA synthetases (Fig. 4; Additional File 2, Table S4). Regarding protein degradation, almost all subunits of the ubiquitin proteasome system (UPS) were present (Fig. 5) as well as autophagy genes, heat shock proteins, and the transcription factor *xbp1. Xbp1* is involved in the ‘unfolded protein response’ (UPR) and in the ER-associated protein degradation (ERAD) pathway [40,41].

Additionally, *BRCA1* was also present in the old subnetwork. This gene is involved in oxidative stress response, and in the transcriptional activation of proteasomal genes by stabilizing the transcription factor *cnc*/*nrf-2* (*cap-n-collar/nuclear factor erythroid 2–related factor 2*) [42]. Other genes in the old subnetwork involved in oxidative stress response and transcriptionally activated by *nrf-2* were thioredoxin and S-glutathione transferase.

##### Other signatures

Additionally, the old subnetwork was characterized by a signature of ecdysone biosynthesis with *ecdysone receptor* (*EcR*), *ecdysone-induced protein 75B* (*Eip75B*), *phantom* and *disembodied*. The presence of *Phosphotidylinositol 3 kinase 68D (Pi3K92E)* links to the IIS pathway.

### Locating age-related co-expression modules in the age-related subnetworks

Finally, we inspected those WGCNA modules with a large fraction of transcripts in the young and old subnetworks as well as those modules that were significantly associated with age. In the young subnetwork, WGCNA modules with a large fraction of transcripts were ‘saddlebrown’ and ‘skyblue4’ which both did not significantly correlate with age. Significantly age-correlated co-expression modules were firebrick2, indianred1 and seashell4 (Additional File 1, Figure S2). No GO terms were significantly enriched for any of these modules.

In the old subnetwork, the modules with a large fraction of transcripts were ‘green’ and ‘paleturquoise’ which again did no significantly correlate with age. The old subnetwork contained transcripts of 13 significantly age-correlated co-expression modules (Additional File 1, Figure S3).

The GO enrichment analysis of these modules revealed several terms involved in protein related functions, including ribosome biogenesis, rRNA processing, protein folding, translation, unfolded protein binding, protein catabolic process, protein transport, tRNA aminoacylation for protein translation, and proteasome core complex (Additional File 1, Figure S4, S9, S10 and Additional File 4, Table S5).

## Discussion

Our study revealed a median maximum reproductive longevity of *C. secundus* queens of 12 years with a maximum lifespan of 13 years when excluding all causes of extrinsic mortality in the laboratory. The curve suggests a rather sudden decline in live expectancy after an age of around 11 years; out of eight queens with a potential age ≥ 12 years, all had died, except one 13-year old queen (Fig. 1). The survival curve indicates a type I survivorship, after queens have successfully founded a colony and without extrinsic mortality.

Our transcriptome study identified six clusters of transcripts that were significantly differentially expressed with age (DETs) (Fig. 2): one cluster for young queens (cluster 2), two for medium aged queens (cluster 3 and 5) and three for old queens (cluster 1, 4, and 6). This implies that three ‘molecular’ life stages can be distinguished in *C. secundus* queens with the third corresponding to old, aged queens that will probably die soon as no queen reached a lifespan beyond 13 years. The co-expression network analysis, which extracted subnetworks based on age-related DETs, resulted in two main subnetworks, a young and an old subnetwork. This implies that there are two *age-related* ‘molecular’ life stages, as DETs/genes of young and medium ages belonged to the same young subnetwork.

### Young subnetwork

The young subnetwork contained DETs characteristic for young and medium ages. This shows the similarity in expression of these two age stages. Not unexpectedly, the young subnetwork indicates an upregulation of transcripts linked with fecundity (*e.g., hts, ball*) and of the TOR pathway (*mipp2*) which has been associated with longevity [34]. The upregulation of *Ac76E* may imply that the IIS pathway is down because this gene is activated by the transcription factor foxo which is inhibited by an upregulated IIS pathway. However, other evidence suggests that, like in other social insects (*e.g.* [12] and references therein), queens are characterized by an upregulated IIS pathway [23]. Additionally, we detected several upregulated TEs-related transcripts associated with signs of an upregulation of the endo-siRNA pathway (*e.g. ago2*, *ars*) which is a transcriptional and post-transcriptional TE-defence mechanism of the soma [32,33,43] (Kim, Lee and Carthew, 2007; Sabin *et al.*, 2009; Piatek and Werner, 2014).

### Old subnetwork

The old subnetwork contained many more transcripts (1,098 vs. 164 in the young subnetwork). Our results imply a physiological stage of upheaval shortly before queens die. There were strong signs of decline and repair at the upstream transcriptional level of epigenetic control as well as at the posttranscriptional level of TE-activity and the proteostasis network.

#### Epigenetic modifications

An upregulation of genes modifying histone marks implied considerable epigenetic changes that lead to altered gene expression as is typical for aging organisms:

First, our results indicate dynamic changes of ‘active’ histone marks of euchromatin because many genes related to H3K4 and H3K36 de/methylation and H4K16 de/acetylation were present in the old subnetwork (Additional File 2, Table S4). For instance, the Tip60-as well as the msl-complex were well represented. Both complexes are involved in acetylation of histones, including H4K16, which, for instance, is indicative for replicative aging in yeast [44]. The upregulation of *Sirt1* may function as an antagonist as it deacetylates H4K16. The same applies for the deacetylases HDAC6 and HDAC3, which can also deacetylase histones [39].

Second, there is also evidence for changes of repressive histone mark of heterochromatin (*e.g.* linked to H3K9 and H3K27 acetylation) (Additional File 2, Table S4). For instance, the old age transcript ALP1 is an antagonist of HP1, the latter is involved in the maintenance of heterochromatin. HP1 generally decreases with age, and its overexpression can lead to an increased lifespan in *D. melanogaster* [45,46].

#### TE activity

In line with a deregulation of repressive histone mark of heterochromatin, several TE-related transcripts were also connected with the old subnetwork (Additional File 2, Table S4). TEs often are accumulated in heterochromatin which silences their activity [47]. Yet, dysregulation of heterochromatin can allow them to become active and this has been associated with aging [48,49]. Also the upregulation of several genes from two TE-defense pathways - the endo-siRNA pathway (*e.g. dicer2*, *Hsc-70-4*, *trsn*) as well as the of piRNA pathways (*e.g.*, piRNA biosynthesis. *armi*, *gasz*, *Hel25E*, *Rm62*; ping-pong cycle: *Tudor-SN*, *qin*) - support the notion of active TEs. Both pathways silence TEs posttranscriptionally [50–52]. TE activity and especially the piRNA pathway has also been associated with aging and the longevity of termite reproductives in another termite species [22].

#### Loss of proteostasis

Our results revealed a very strong proteostasis signal indicative for an upheaval in protein synthesis, protein folding and protein degradation in old queens. Many genes from the proteostasis network were detected in the old subnetwork (Additional File 2, Table S4). They indicate an upregulation of protein degradation, together with protein synthesis and protein folding (Figs. 4 and 5). This is unusual because old organisms are typically characterized by a downregulation of all these processes (Fig. 6).

**Figure 6.**
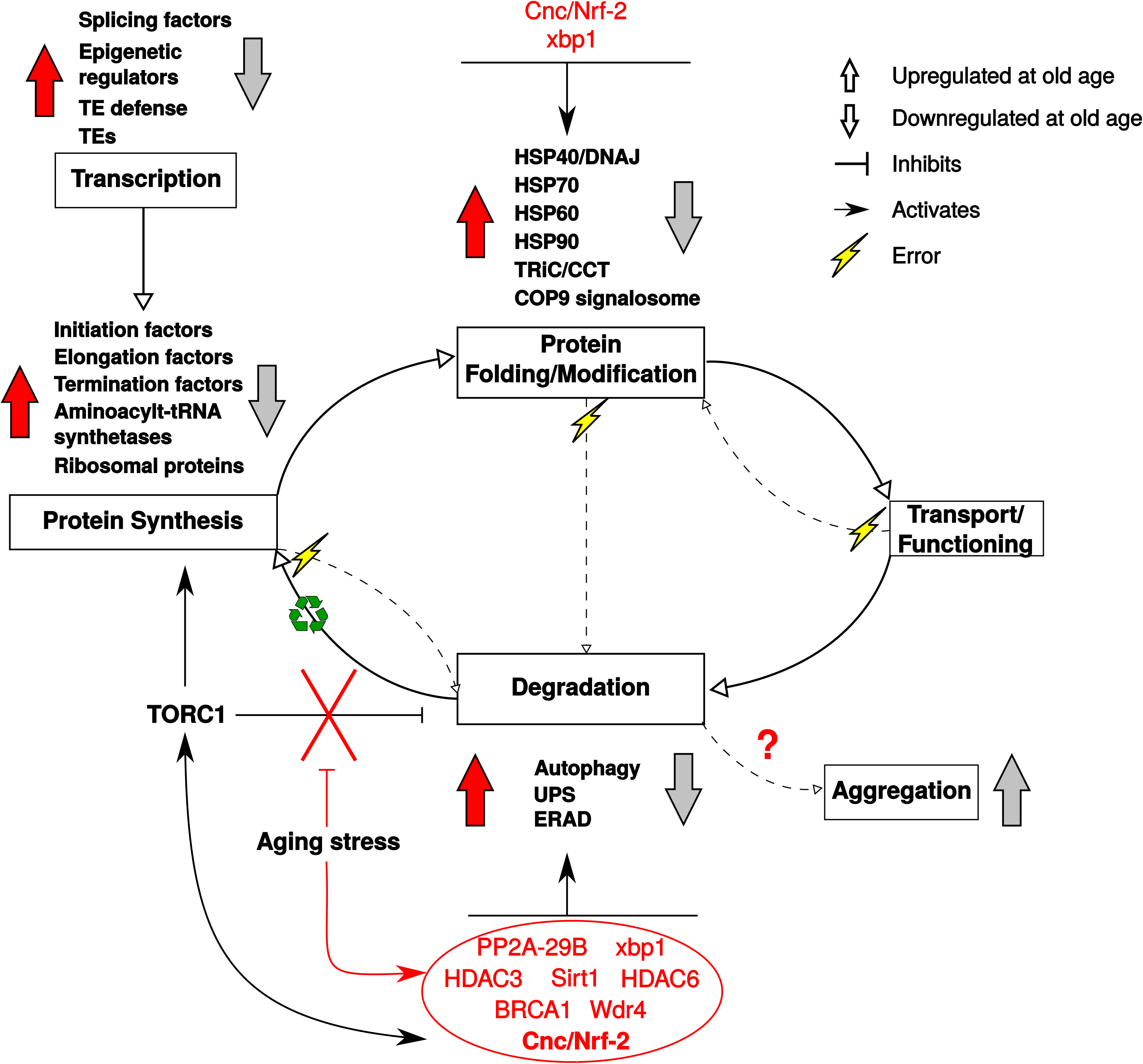
Transcription- and proteostasis-related expression pattern in old *Cryptotermes secundus* queens. Depicted is the expression patterns of genes related to transcription and proteostasis for old *C. secundus* queens (red arrows) in contrast to that reported for other species (grey arrows). Old *C. secundus* queens were characterized by a very strong proteostasis signal indicative of an upregulation of protein degradation, together with protein synthesis and protein folding. This is unusual because old organisms are typically characterized by a downregulation of these processes. The simultaneous activation of protein synthesis and degradation in old *C. secundus* queens can be explained by the activity of the transcription factor cnc/Nrf-2 (for more details, see text). The inner cycle arrows depict the protein life cycle; dashed arrows indicate the special case when mistakes/ errors occur. After a protein is degraded, its components are recycled.

##### Protein synthesis

Many transcripts coding for ribosomal proteins and aminoacyl-tRNA synthetases were found in the old subnetwork, indicative of upregulated protein synthesis (Additional File 2, Table S4). This is further supported by a strong signal of an active TORC1 system which promotes protein synthesis (Fig. 7 [53–55]). Thus, for instance, many downstream eukaryotic initiation factors (eIF4A, eIF4B, eIF4E, eIF4G) and eukaryotic elongation factor (eEF2) transcripts were found in the old subnetwork (Fig. 4). They activate, ribosome biogenesis, translational elongation, and cap-dependent translocation (Fig. 7).

**Figure 7.**
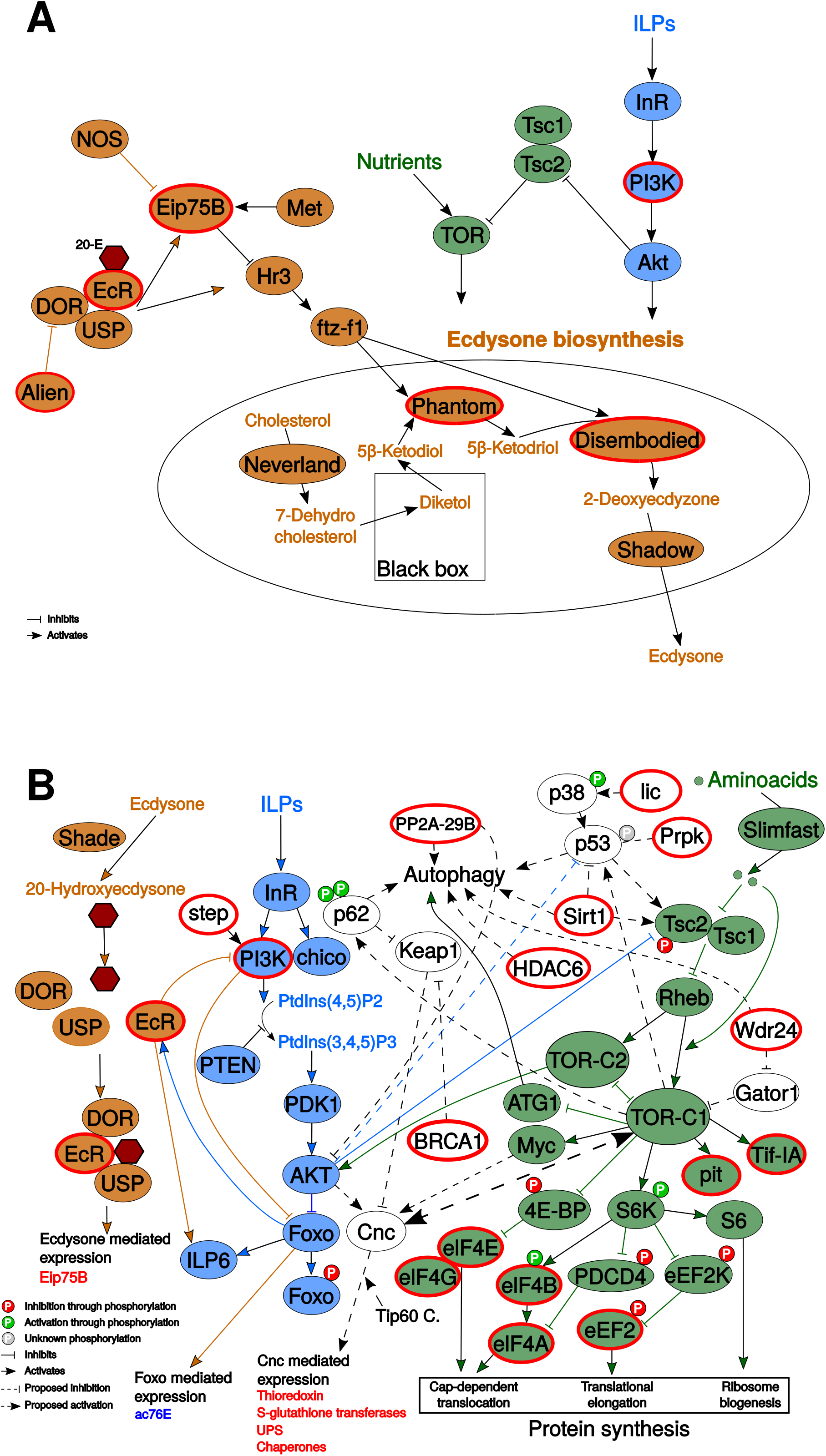
Aging signal of C. secundus queens in relation to known aging pathways. Shown are simplified IIS (insulin/insulin-like growth factor signaling; blue), TOR (target of rapamycin; green), and ecdysone (brown) pathways and their interactions with an emphasis on **A.** ecdysone biosynthesis and **B.** protein synthesis and degradation. Red encircled genes were members of the old subnetwork, and thus highly expressed in old queens. Important genes that regulate protein synthesis and degradation are depicted in white. In short, the TOR pathway controls cell growth and metabolism in response to amino acid availability. It is generally composed of two main complexes: TORC1 and TORC2 [55]. Activation of TORC1 promotes mRNA translation, for instance, via S6K /eIF4B / eIF-4a and 4E-BP / eIF4E. Additionally, active TORC1 inhibits autophagy by targeting upstream components necessary for autophagy activation, like Atg1. TOR interacts with IIS, which also regulates multiple physiological functions, including aging. Generally, an active IIS pathway can activate the TORC1 complex via phosphorylation and inactivation of Tsc2 by AKT. AKT inhibits the transcription factor foxo via phosphorylation, which results in the inhibition of transcription of many downstream genes, e.g. involved in lifespan extension, stress response and autophagy. Stress induced *Cnc* can activate TORC1 in a positive feedback loop (big dashed arrow). It may be responsible for the simultaneous upregulation of protein synthesis and degradation. For more information, see text. Figures are adapted after [53–55].

##### Protein degradation

Normally, an active TORC1 system is associated with a downregulation of protein degradation as it inhibits proteolytic systems [54–56] and autophagy (*e.g.,* upregulated TORC1 inhibits ATG1, which is necessary for autophagy activation; Fig. 7). Surprisingly, however, we found strong evidence of upregulated protein degradation in the old subnetwork. Several transcripts linked to autophagy, almost all subunits of the UPS, the UPR-, and the ERAD pathway as well as heat shock proteins characterized the old subnetwork (Figs. 5 and 6; Additional File 2, Table S4).

##### Linking protein synthesis and degradation

The simultaneous upregulation of protein synthesis and autophagy might be explained by a stress response. In *D. melanogaster* as well as in humans, under stress conditions upregulated TORC1 enhances an oxidative stress response, controlled by the transcription factor *cnc*, a homolog of human *nrf* genes (Fig. 7). Ubiquitinated proteins and damaged mitochondria activate *cnc/ nrf-2* via p62, supported by upregulated TORC1, which then activates oxidative stress response genes [57,58] (Ichimura *et al.*, 2013; Aramburu *et al.*, 2014). Additionally, *cnc* is known to activate chaperones (protein folding) and the proteasome [59], and this has been associated with lifespan expansion in *D. melanogaster* and *Caenorhabdtis elegans* [60,61].

Support for the conclusion of a stress-associated, active *cnc* transcription factor in old queens comes from several transcripts in the old subnetwork (Fig. 7): (i) BRCA1, which indirectly actives *cnc* by inhibiting *cnc* inhibitor *Keap1*, and (ii) the *Tip 60* complex as well as genes that are transcriptionally activated by *cnc*, such as *thioredoxin, S-Glutathione transferases* and *UPS* genes (Fig. 7).

Hence, our results imply that old *C. secundus* queens are in a stage of stress. They have mounted stress response systems mediated by *cnc*, including protein degradation and protein folding. However, it is unusual that old queens can do this. In *D. melanogaster*, only young individuals can mount this stress response [59] (Tsakiri *et al.*, 2013). The constant activation of the proteasome in these very old queens may lead to their death (note, the studied queens had reached their maximum lifespan, we never had older queens) as the constant activation of the proteasome in transgenic flies was detrimental for survival [59].

### Comparison with other social insects

There has been a debate about the role of oxidative stress to explain the long lifespan of social Hymenoptera queens; yet evidence is inconclusive (*e.g.* reviewed by [19–21]). For instance, markers of oxidative stress in the brain of honey bee workers do not increase with age, although they live shorter than queens [62]. In the ant *Lasius niger,* both workers and males show higher activity of the antioxidant superoxide dismutase than queens, but both live shorter than queens [63]. These studies have shown that higher expression of oxidative stress response genes do not necessarily correlate with longer lifespans. For the termite *Reticulitermes speratus*, it has been suggested that queens are better protected against oxidative stress as qRT-PCRs studies showed a higher expression of the antioxidants catalases and peroxiredoxins in queens compared to workers [17], while kings were characterized by a high expression of *BRCA1* (in the fat body) compared to workers [15]. Unfortunately, the age of the studied reproductives is not known. It would be of interest to study expression of these genes with age, as this would contribute to a better understanding of the aging process but such studies are rare.

## Conclusion

Our results imply that *C. secundus* queens do not age gradually, rather at old age there is a physiological stage of upheaval, characterized by signs of stress (activity of TEs, active *crc*) and defense (piRNA pathways) / repair (protein degradation and synthesis) before the animals die. This apparently sudden decline is in line with the few life history records of social insect queens that exist. They also found no signs of gradual senescence but an abrupt death (*e.g.,* the ant *Cardiocondyla obscurior* [64]; see also Fig. 1). This stresses the importance of using queens of known age for aging studies as processes revealed from middle-aged versus old queens probably differ considerably. Our study is the first addressing the aging process for a social insect by studying the complete lifespan of queens.

## Methods

Figure S1 (Additional File 1) provides a schematized workflow of the analyses described in the following sections.

### Collection and colony maintenance

From 2002 until 2016, *C. secundus* colonies were collected from mangroves near Palmerston - Channel Island (12°30’ S, 131°00’ E; Northern Territory, Australia) when they were less than one year old [65]. Colonies of an age of less than one year can be identified by the size and slightly lighter sclerotization of the founding reproductives (primary reproductives), the presence of less than 20 workers and short tunnel systems of a few centimeters. All collected colonies were transferred to *Pinus radiata* wood blocks and transported to the laboratory in Germany, where they were kept under standardized conditions in a climate room with a temperature of 28 °C, 70% humidity and a 12h day/night rhythm (Additional File 2, Table S1). Under these conditions, colonies develop like in the field (see [66]). Up until 2019 wood blocks were split and the colonies were extracted to determine survival of reproductives.

### Survival analysis

The survival of primary queens (and kings) was determined by their presence /absence. Founding (primary) queens can be identified by their dark brown color, compound eyes and wing abscission scars. If the primary reproductives had died and the colony was still alive, they had been replaced by neotenic replacement reproductives which lack these traits. The median longevity of queens was determined using Kaplan Meier survival analysis in SPSS 23 [67]. Only colonies that survived the transfer from the field to the laboratory in Germany and the re-establishment in the new wood block were used for this analysis. This resulted in a sample size of 41 colonies. Overall, we had surviving primary queens from an age of one year up to a maximum of 13 years. The three potentially 14- and 15-year old queens were all dead.

### Transcriptome study

#### RNA extraction and sequencing

RNA was extracted from twelve queens with different chronological age since onset of reproduction from two years until 13 years: 2, 3, 4, 5, 6, 7, 8, 9, 10 (two samples), 11, and 13 years. In colonies older than 13 years, the original queen was always replaced by a neotenic replacement queen.

An in-house protocol was followed for RNA extraction (see [23]). Individuals were placed on ice and the gut was removed and discarded. The head together with the thorax were used for RNA extraction. Samples were transferred into peqGOLD Tri Fast™ (PEQLAB) and homogenized in a Tissue Lyser II (QIAGEN). Chloroform was used for protein precipitation. From the aqueous phase, RNA was precipitated using Ambion isopropyl alcohol and then washed with 75% ethanol. Obtained pellets were solved in nuclease-free water. DNA was subsequently digested using the DNase I Amplification Grade kit (Sigma Aldrich, Cat. No. AMPD1). We performed an RNA Integrity Number Analysis (RIN Analysis) measuring the RNA concentration with the Agilent RNA 6000 Nano Kit using an Agilent 2100 Bioanalyzer (Agilent Technologies) for quality control. Samples with total RNA were sent on dry ice to Beijing Genomics Institute (BGI) Tech Solutions (HONGKONG) Co. and then to the BGI-Shenzhen (PR China) for sequencing. The preparation of the cDNA libraries was performed by BGI according to their internal and proprietary standard operating procedure. The cDNA libraries were paired-end sequenced (not-strand specific) on Illumina HiSeq 2500 and 4000 platforms (100 base pairs read length and about 4 Giga bases per sample). Index sequences from the machine reads were demultiplexed and a quality -check and filtering of raw reads was done using the package soapuke (-n 0.05 -l 20 -q 0.2 -p 1 -i -Q 2 -G -- seqType 1 and -A 0.5, http://soap.genomics.org.cn/).

#### Processing of RNASeq raw reads

FastQC (v0.11.4) [68] was used to evaluate the quality of the cleaned raw reads. To obtain a transcript count table, the cleaned raw reads were pseudo-aligned with Kallisto (default settings, v0.43.0) [69] (Bray *et al.*, 2016) against a *C. secundus* transcriptome obtained from a draft version of the *C. secundus* genome (with estimated gene and transcript models, see [23,70]. The counts estimated by Kallisto were normalized using DESeq2 (v1.18.1, count/size factor) [71] in R (v3.4.4) [72].

#### Time course analysis to identify age-associated differentially expressed transcripts (DETs)

The normalized counts were used as input for the R package Iso-MaSigPro (v1.50.0) [73] to test for differentially expressed transcripts (DETs) across time. Iso-MaSigPro is designed for the analysis of multiple time course transcriptome data. It implements negative binomial generalized linear models (GLMs) [73,74]. Significantly differentially expressed transcripts (FDR corrected p-value set to 0.05) were clustered using the clustering algorithm mclust in Iso-MaSigPro [73] resulting into six Iso-MaSigPro clusters (Additional File 2, Table S2).

#### Weighted Gene Co-expression Network Analysis (WGCNA) to identify networks of co-expressed transcripts

To obtain networks of co-expressed transcripts that were categorized as modules we performed a Weighted Gene Co-expression Network Analysis (WGCNA). The counts obtained with Kallisto (v0.43.0) [69] were transformed using variance stabilizing transformation (vst) as implemented in DESeq2 (v1.18.1) [71]. The vst transformed counts were used to perform a co-expression network analysis with the R package WGCNA (v1.63) [75]. For more details on the methodology, see [75–77]. In short, (Additional File 1, Figure S1, workflow, right side), a similarity matrix was built by calculating Pearson correlations between the expression values of all pairs of transcripts. Using the similarity matrix, a signed weighted adjacency matrix was obtained as described by the formula:

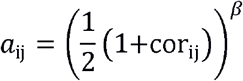

Where cor_ij_ is the Pearson correlation between the expression pattern of transcript ‘i’ and transcript ‘j’ (the similarity value). The value of β was chosen based on the soft-thresholding approach [75]. With this value of β, we obtained a weighted network with an approximate scale-free topology (β=14, scale-free topology R^2^ = 0.84). In a signed weighted adjacency matrix negative and small positive correlations get negligibly small adjacency values shifting the focus on strong positive correlations. Seeing the adjacency matrix as a network, the nodes correspond to the transcripts and the connections between nodes correspond to the adjacency values (transformed correlation coefficients). A topological overlap matrix (TOM), which in addition to the adjacency matrix considers topological similarity (shared neighbors reinforce the connection strength between two nodes), was constructed using the adjacency matrix [78]. To define transcript modules, a hierarchical clustering tree was constructed using the dissimilarity measure (1-TOM). Transcript modules were defined by cutting the branches of the tree using the Dynamic Hybrid Tree Cut algorithm [79] and the minimum module size was set to 30 transcripts. Transcript modules with similar expression profile were merged by hierarchical clustering of the eigengene correlation values. Briefly, a hierarchical clustering tree was created with the eigengene dissimilarity measure (1-correlation coefficient of eigengenes) and a tree height cut of 0.20 was used (corresponds to a eigengene cor ≥ 0.80). Eigengenes were calculated with the function moduleEigengenes (default settings) [75]. A module eigengene corresponds to the first principal component of the module and can be seen as a weighted average expression profile [75]. To find significantly associated modules with age, correlations between age and eigengenes of the merged modules were calculated. Each module was named after a color by WGCNA.

The adjacency matrix of the WGCNA was visualized using Cytoscape (v3.7.1) [80], only including pairs of nodes with a cor_ij_ ≥ 0.90. The color of each module corresponds with the respective module name (*e.g.*, saddlebrown color for the saddlebrown module).

To identify co-expression modules containing age-related DETs, we looked for age-related DETs from the Iso-MaSigPro analysis in the WGCNA modules. Those modules which were significantly correlated with age and which contained the highest number of Iso-MaSigPro DETs were further inspected.

### Identifying/Extracting age-related subnetworks based on age-related DETs

To identify age-related subnetworks within the co-expression network, we combined the results of the Iso-MaSigPro analysis with those from the WGCNA and extracted subnetworks that were based on age-related Iso-MaSigPro DETs. Therefore, we extracted 1^st^ and 2^nd^ neighbors of DETs based on the WGCNA co-expression network (*i.e.,* the visual representation of the adjacency matrix). To do this, we used the ‘First neighbors’ function of Cytoscape. We selected an age-related DET from Iso-MaSigPro as transcript of interest. By calling the function, the neighboring transcripts were selected, which were then extracted to form a subnetwork. By calling the function twice, one obtains the one- and two-step neighbors (called ‘second level neighborhood’) of the transcript of interest. This was done for each DET identified in IsoMaSigPro.

The obtained subnetworks were clearly separated in those containing young Iso-MaSigPro DETs (young subnetworks) and those containing old Iso-MaSigPro DETs (old subnetworks). The largest subnetwork obtained for each group was used for further analysis paying attention to both, transcript identity as well as WGCNA module content. Thus, we looked, for instance, for WGCNA modules that had been identified to be age-related within the global WGCNA in these subnetworks. The AutoAnnotate Cytoscape plug-in (v1.3) [81] was used to annotate the subnetworks using the clustering algorithm ‘Markov Cluster’ (MCL) [82] to define and visualize sub-clusters, and the labeling algorithm ‘Adjacent Words’ to label the sub-clusters. The Cytoscape plug-in BiNGO (v3.0.3) [83] was used for GO enrichment analysis. The p-values of the GO enrichment analysis were adjusted for multiple testing using the FDR approach [84]. Subnetworks were graphically processed with Inkscape (v0.91, www.inkscape.org).

### Transcript (functional) annotation

Nucleotide and protein sequences were obtained from the draft version of *C. secundus* genome [23,70]. For annotation, the translated transcripts were searched against the Pfam database (Pfam A, release 30) [85] with the software *hmmscan* (option --cut_ga, HMMer v.3.1b2) [86] and against the InterPro database with the software InterProScan (v5.17-56.0) [87]. Additionally, we did a BLASTX search (NCBI BLAST suite v. 2.3.0) [88] with an e-value of 1e^−05^ as cut-off against the protein coding sequences of the termite *Zootermopsis nevadensis* (official gene set version 2.2) [89]. To further assist the annotation, we inferred a set of clusters of orthologous sequence groups (COGs) from the official gene sets at the amino acid level of *C. secundus* (draft version) and *D. melanogaster*, and a BLASTP search of *C. secundus* sequences against the protein coding sequences (longest isoforms only) of *D. melanogaster* with a threshold e-value of 1e^−05^.

To detect possible TEs, transcripts were searched against the Dfam database (v2.0) [90] using nhmmer [91]. A transcript was considered TE related if there was a hit against the Dfam database and the other annotation sources (Pfam, Interpro and BLAST) were not pointing in the direction of a known gene.

### Gene identification and construction of gene trees

In addition to the functional annotation, we inferred phylogenetic trees for selected transcripts (Supplementary Archive 1, DRYAD, doi available upon acceptance of the manuscript). Following the procedure described in [23], we retrieved protein coding sequences of the respective cluster of orthologous sequence groups (COGs) from OrthoDB v.9.1 [92] for the following species: *D. melanogaster* (DMEL), *Apis mellifera* (AMEL), *Cardiocondyla obscurior* (COBS), *Polistes canadensis* (PCAN)*, Tribolium castaneum* (TCAS), *Z. nevadensis* (ZNEV) and *Blattella germanica* (BGER). COGs were identified using text search by searching for the gene name or IDs of *D. melanogaster.* In case Selenocysteine (U) was included in sequences, “U” was replaced by “X” to avoid problems in downstream analyses since many programs cannot handle Selenocysteine properly. Protein sequences of COGs of above listed species were separately aligned with MAFFT (v.7.294b) applying the G-INS-i algorithm and otherwise default options [93]. For each multiple sequence alignment, a profile hidden Markov model (pHMM) was built with hmmbuild (HMMER v.3.1b2) [86]. Then the pHMM was used to search (hmmsearch) for corresponding protein coding sequences in *C. secundus* and *Macrotermes natalensis* to identify orthologous candidate sequences for each COG in both species. For gene (transcript) tree inference, we only kept sequences with a threshold e-value of ≤ 1e^−40^. In addition, we annotated all candidate sequences identified in *C. secundus* and *M. natalensis* against the Pfam database (Pfam A, release 30) using hmmscan (HMMER v.3.1b2).

To infer phylogenetic gene trees, we merged for each COG the COGs (amino-acid level) of the seven species listed above with the putatively orthologous amino-acid sequences of *C. secundus* and *M. natalensis*. We generated multiple sequence alignments for a total of 29 genes of interest applying MAFFT (G-INS-i, see above). Ambiguously aligned sequence sections were identified with Aliscore (v. 2 [94,95]; settings: -r: 10000000 and otherwise defaults) and removed with Alicut (v. 2.3, https://www.zfmk.de/en/research/research-centres-and-groups/utilities; masked alignments provided as Supplementary Archive S1 (deposited at DRYAD, doi available upon acceptance of the manuscript). Phylogenetic trees were inferred with IQ-TREE (1.7-beta12 [96] for each gene. The best model was selected with the implemented ModelFinder [97] from all available nuclear models implemented in IQ-TREE plus the two protein mixture models LG4M and LG4X [98] based on the Bayesian Information Criterion (BIC). We applied default settings for rates and the number of rate categories. Statistical support was inferred from 2,000 non-parametric bootstrap replicates. Unrooted trees with the bootstrap support mapped were visualized with Seaview (v4.5.4 [99]) and provided in Newick Format with Supplementary Archive S1 at DRYAD (doi available upon acceptance of the manuscript).

## Ethics approval and consent to participate

Not applicable

## Consent for publication

Not applicable

## Competing interests

The authors declare that they have no competing interests.

## Availability of data and materials

Raw sequence reads are deposited on NCBI (Bioproject XXX, BioSample Accessions see Additional File 2, Table S1 – Bioproject numbers and BioSample Accessions will be provided when obtained from NCBI). Additional supplementary data are deposited on the Dryad digital repository DRYAD (DOI provided upon acceptance of the final manuscript). For Supplementary Material please contact

## Funding

This research was supported by the Deutsche Forschungsgemeinschaft by grants to JK (DFG; KO1895/16-1; KO1895/20-1), one within the Research Unit FOR2281.

## Contributions

JK designed the study, JK and MMK collected and identified the termite samples, MMK performed all transcriptome analyses, KM helped with gene identification, with data processing and inferred gene trees, JK did survival analysis and supervised the study, all authors wrote the paper.

## Acknowledgments

We thank Florentine Schaub for assistance in the field and wet lab, Daniela Schnaiter for termite colony maintenance, Charles Darwin University (Australia), and especially S. Garnett and the Horticulture and Aquaculture team, provided logistic support to collect *C. secundus*. The Parks and Wildlife Commission, Northern Territory, the Department of the Environment, Water, Heritage and the Arts gave permission to collect. Permit numbers 2002 until 2016, export permit numbers: PWS2001_1508, PWS2003_39852, PWS2004_5769, PWS2007_4154, PWS2010_6997, PWS2014_001342, PWS2016_001559; collection permit numbers: 15656, 18310, 26851, 30073, 36401, 51402, 59044. The study was conducted according to the Nagoya protocol. KM thanks Ondrej Hlinka and the CSIRO IM&T HPC Cluster team.

## Additional Files

### Additional File 1 (Additional_File_1.pdf): Supplementary Figures S1-S10

Figure S1: Schematic workflow. Details are explained in the Methods.

Figure S2: WGCNA modules of co-expressed transcripts that negatively correlate with age. Modules marked with an asterisk were found in the young subnetwork. Modules are named after colors by WGCNA.

Figure S3: WGCNA modules of co-expressed transcripts that positively correlate with age. Modules marked with an asterisk were found in the old subnetwork. Modules are named after colors by WGCNA.

Figure S4: GO enrichment for the WGCNA module ‘tan’ that positively correlated with age and which had many old-age DETs. Details are shown for BP (Biological Process), which revealed an enrichment of transcripts for ribosomal and tRNA related functions.

Figure S5: Young and old transcript subnetworks corresponding to the second level neighborhood of Iso-MaSigPro DETs. Age-related DETs were located in the WGCNA co-expression network and these DETs and their one- and two-step neighbors (*i.e.,* the ‘second level neighborhood’) were then extracted from the co-expression network to provide the shown networks.

Figure S6: BiNGO GO enrichment (Biological Process) for the young subnetwork. No terms were significantly enriched after correcting for multiple testing (FDR).

Figure S7: BiNGO GO enrichment (Biological Process) for the old subnetwork. Colored nodes are GO terms that were significantly enriched after correcting for multiple testing (FDR).

Figure S8: BiNGO GO enrichment of (Molecular Function; Cellular Component) for the old subnetwork. Colored nodes are GO terms that were significantly enriched after correcting for multiple testing (FDR).

Figure S9: BiNGO GO enrichment (Biological Process; Molecular Function; Cellular Component) for the ‘green’ WGCNA module, which is part of the old subnetwork. Colored nodes are GO terms that were significantly enriched after correcting for multiple testing (FDR).

Figure S10: BiNGO GO enrichment (Biological Process, Molecular Function; Cellular Component) for the ‘paleturquoise’ WGCNA module, which is part of the old subnetwork. Nodes in color are GO terms significantly enriched after correcting for multiple testing (FDR).

### Additional File 2 (Additional_File_2.xlsx): Supplementary Tables S1-S4

Table S1: Sample information of samples included in this study.

Table S2: Differentially expressed transcripts of the Iso-MaSigPro analysis, separately for cluster 1-6.

Table S3. Transcripts in the young subnetwork (SNW).

Transcripts in the young subnetwork (SNW) classified into major categories; TE-related transcripts of the young subnetwork (SNW).

Table S4. Transcripts in the old subnetwork (SNW).

Transcripts in the old subnetwork (SNW) classified into major categories; TE-related transcripts of the old subnetwork (SNW).

### Additional File 3 (Additional_File_3.pdf)

WGCNA module-age associations. Listed are the eigengene values for each module.

### Additional File 4 (Additional_File_4.xlsx): Supplementary Table S5

Go terms for enriched differentially expressed transcripts (DETs) included in BiNGO module green; BiNGO module paleturquoise; BiNGO module tan; BiNGO module Thistle2; BiNGO module snow; BiNGO module cyan; BiNGO module deeppink1; BiNGO module navajowhite; BiNGO module blue2_NS; BiNGO module cornflowerblue_NS; BiNGO module pink3_NS; BiNGO module steelblue4_NS.

### Supplementary Archive 1

The Supplementary Archive includes i) masked multiple sequence alignments (MSAs) of 29 selected genes (subdirectory “MSAs” used for ML gene tree inference in FASTA format) in IQ-TREE and ii) inferred ML gene trees (subdirectory “ML_gene_trees” in NEWICK format) for each of the 29 genes including non-parametric statistical bootstrap support. These gene trees additionally helped to ensure a proper assignment of transcripts to repective genes.

## Notes

### Competing Interest Statement

The authors have declared no competing interest.

